# Enrichment of bioplastic degraders in mesophilic compost from widespread degradation potential in the environment

**DOI:** 10.64898/2025.12.16.694645

**Authors:** Tan Suet May Amelia, Shu Yuan Yang

## Abstract

Polylactic acid (PLA) is a polymer that is known to exhibit compostability at thermophilic temperatures, and this activity is thought to be connected to the presence of PLA hydrolyzability in environmental microbes. We recently developed a set of compost that can biodegrade PLA at mesophilic conditions, and one possible reason underlying our success could be due to the enrichment of PLA hydrolyzability. Here, we investigated the potential selection of bioactivities related to PLA breakdown in our trained compost and surveyed the occurrences of those activities in the environment for comparison. Ten different environments were sampled, including PLA, larval gut of black soldier flies, and organic material from our trained compost, as well as terrestrial soil, estuarine sediment, brackish water, shell biofilm, coastal stranded polystyrene, bottle, and bottle cap. We found a small fraction of cultivable bacteria in many samples that harbored PLA degradability. For our trained compost, PLA-degrading isolates were twice as efficient as those from other environments, even though the frequency at which they were detected was not significantly higher. These findings suggested that PLA breakdown ability is commonly present at a low percentage in most environments, and that our trained compost has been able to select for more effective isolates. As this enhancement is likely insufficient to explain the increase in PLA compostability in our trained compost compared to standard mesophilic composts, we propose that additional microbial activities are needed to act synergistically and overcome the requirement for elevated temperature in PLA composting.

**IMPORTANCE:** This study represents our work in investigating the biodegradation activity of the most common bioplastic, PLA, in the environment and in a special compost we recently developed that exhibited the novel ability of being able to achieve PLA composting at ambient temperatures. Our work is a rare survey that compares PLA hydrolytic activity across different environments, helping unmask the underlying prevalence of environmental PLA hydrolysis activity, as well as whether our special compost is especially enriched for such activity, would facilitate the design of PLA biodegradation implementation strategies. We found that PLA hydrolytic activity was generally present in environmental microbes at low frequencies, and that our special compost selected for those that were more efficient. However, full PLA compostability under mesophilic conditions likely depends on embedded, synergistic microbial functions beyond hydrolysis alone, motivating future work to disentangle complementary activities that collectively enable complete breakdown.

## INTRODUCTION

Bioplastics are promoted for their biodegradability potential and are therefore widely used in single-use products, yet this dependence reduces their intended environmental benefits, as minimal biodegradation of bioplastic waste has been implemented (1, 2). Polylactic acid (PLA) is the leading commercial bioplastic, representing global bioplastics production of about 37% in 2024 to a projection of 42% in 2029 (3, 4). PLA is widely recognized as industrially compostable (5). However, industrial composting of PLA consists of a prolonged abiotic degradation phase that requires high temperatures (58°C and above) and extended processing periods; thus, incomplete disintegration, increased processing costs, and the accumulation of recalcitrant polymer residues in industrial facilities are common problems encountered that curb enthusiasm for such practices (6, 7).

Mesophilic composting (typically 30-45°C) offers a simpler, low-cost option achievable in small-scale settings (8, 9), but biodegradation of PLA is generally negligible under these conditions in spite of multiple PLA depolymerization enzymes (mainly hydrolytic enzymes) active at mesophilic temperature ranges being reported (10, 11). Material modification (e.g., blending with alternative polymers), biostimulation (e.g., introducing biostimulants or catalysts), and bioaugmentation (e.g., introducing microbial agents) techniques have been shown to enhance degradation, indicating progressing research in the mesophilic composting of PLA (9, 12, 13).

In our recent work, a specially trained compost that acquired mesophilic PLA biodegradability has been reported (14). Many microorganisms that break down PLA have been reported, but this potential has scarcely been translated into practical, large-scale PLA waste treatment (15). The incomplete understanding of mesophilic PLA-degrading microbes and their mechanisms necessitates investigations on the prospects of mesophilic compost as a microbial reservoir for PLA degraders. Therefore, we quantified and compared the viability and clearing ability of mesophilic isolates on PLA from our trained compost and diverse matrices, enabling us to probe the prevalence of PLA hydrolytic capacity and pinpoint environments that may be enriched. Specifically, we evaluated isolates from bulk compost, compost-derived PLA fragments, compost-derived larval gut, terrestrial soil, estuarine sediment, brackish water, shell biofilm, and coastal stranded plastics.

## MATERIALS AND METHODS

### Media preparation and isolate maintenance

Four types of mineral salt medium (MSM) agar plates supplemented with PLA, polyhydroxyalkanoate (PHA), polystyrene (PS), and polyethylene terephthalate (PET) polymers were prepared. The four types of plastic powder-containing agar plates used consisted of a base of MSM: 1.50 g/L K_2_HPO_4_, 2.08 g/L Na_2_HPO_4_, 1 g/L NH_4_Cl, 0.1% (v/v) of 0.1M MgSO_4_·7H_2_O, 0.1% (v/v) of trace elements, 14 g/L agar, and the respective polymers (3 g/L for PLA and PHA; 1.5 g/L for PS and PET) (16). PLA (25 µm particles, Unic Technology) and PHA (30 µm particles, Unic Technology) available readily in powder form were used directly to make PLA- and PHA-MSM agar media, while emulsified PS (5 mm pellets, Chimei) and emulsified PET (5 mm pellets, Huixin) were formed in advance by double emulsion solvent evaporation described below prior agar preparation. The trace element solution consisted of 2.78 g/L FeSO_4_·7H_2_O, 1.98 g/L MnCl_2_·4H_2_O, 2.81 g/L CoSO_4_·7H_2_O, 1.67 g/L CaCl_2_·2H_2_O, 0.17 g/L CuCl_2_·2H_2_O, and 0.29 g/L ZnSO_4_·7H_2_O (17).

Emulsified PLA-MSM agar components matched PLA-MSM agar, except PLA powder was substituted by emulsified PLA fine powder (1.5 g/L) obtained by double emulsion solvent evaporation to achieve homogenous PLA-MSM. 1:1 of 7% (w/v) PLA (25 µm, Unic Technology) in dichloromethane (DCM) and 0.5% (w/v) polyvinyl alcohol (PVA) aqueous solution were homogenized at 30,000 rpm for 20 s. The emulsion was stirred for 48 h until >50% evaporated to gradually remove DCM, then centrifuged at 10,000 rpm for 5 min. The supernatant was discarded, and the pellet was rinsed with distilled water three times to remove residual PVA. Each plate received 20 mL emulsified PLA-MSM to standardize the thickness. Emulsified PS was prepared identically, while emulsified PET used phenol (70°C, 30 min) instead of DCM to dissolve PET (18).

Tween 80-MSM (TW80-MSM) agar consisted of 1% (v/v) of Tween 80 in MSM and validated with emulsification index assay (E24, Figure 5) (19). Congo red agar (CRA) consisted of 30 g/L tryptic soy broth (TSB), 20 g/L sucrose, 15 g/L agar, and 0.8 g/L of Congo red dye (20). Nutrient agar (NA) comprised 3 g/L meat extract, 5 g/L peptone, 5 g/L NaCl, and 15 g/L agar. Nutrient-rich (NR) broth comprised of 10 g/L peptone, 10 g/L meat extract, and 2 g/L yeast extract.

### Collection of samples

All environmental samples were collected using sterile equipment and transported to the laboratory in an insulated container within 2 h for immediate processing. From our trained compost (14), PLA pieces, brown compost material not in contact with PLA, and black soldier fly larvae were collected, with the latter aseptically dissected to obtain gut tissue and associated microbes. Three coastal plastic waste types, bottle cap (polyethylene, PE), bottle (PET), and Styrofoam (extruded form of PS), were sampled from Changtanli Fishing Harbor, Keelung (25.1412° N, 121.8018° E). Three marine environmental samples (sediment, brackish water, shell biofilm) were collected near Lishui Fishing Harbor, Wuxi, Taichung (24.2014° N, 120.4913° E). Terrestrial soil was sampled at Chang Gung University, Taoyuan, Taiwan (25.0334° N, 121.3898° E).

### Bacterial strain isolation

Water samples were directly serially diluted for isolation and screening. Plastic and larval gut samples were cut into smaller pieces (approximately 2-6 mm^3^) using sterile scissors before weighing and gently shaken for 3 s in sterile PBS to remove loosely attached bacteria prior to vortexing in PBS-filled microcentrifuge tubes. 0.3 g of the samples were then added to 1.0 mL sterile PBS, vortexed for 10 s, sonicated at room temperature for 15 min, vortexed for 10 s again, then rested for 10 min to allow sedimentation of turbid samples before serial dilution up to x10^-6^. Next, aliquots of 0.1 mL were spread on NA or PLA-MSM plates, then incubated for 96 h at 30°C (21, 22). Distinctness determinations of isolates were made by a combination of criteria, including cellular and colony morphology, and a subset of isolates were sequenced for their 16S genes to validate the classification results. All experiments were performed in triplicate.

### Bacterial strain screening

Non-selective isolation and screening were performed by streaking all distinct isolates onto agar plates containing different types of plastic, and the numbers of cultivable isolates were used for calculating cultivability hit rates by the following equation:

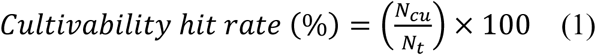

in which *N_cu_* is the number of cultivable isolates after screening and *N_t_* is the number of total isolates before screening.

The numbers of isolates that can generate clearance zones on various plastic-containing plates were also tallied, and clearability hit rates were then determined with this equation:

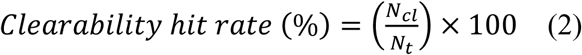

in which *N_cl_* is the number of clearance-causing isolates after screening and *N_t_* is the number of total isolates before screening.

Selective isolation and screening were performed by plating microbial samples directly onto PLA-MSM plates without first being plated on NA. The number of distinct isolates, and those that cause clearance zones using this alternative method, were used to calculate selective isolation clearability hit rates.

### Quantitative polylactic acid (PLA) clearance assay

Quantitative analysis of PLA clearing efficiency was performed by preculturing each test isolate in a 10-mL glass test tube containing 3 mL of NR broth for 8 h at 25°C and an agitation speed of 80 rpm. 10 µL of an OD_600_=1.0 culture was then dropped onto the center of an emulsified PLA-MSM plate and incubated at 30°C for 21 days.

Measurements of the diameter of the bacterial spot and the zone of clearance were taken on days 7, 14, and 21, with diameters at four different locations of each clearance zone and colony taken and averaged (23). The values were used for calculating clearance indices using the following equation:

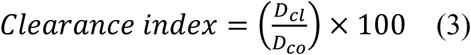

in which *D_cl_* is the diameter of a given clearance zone and *D_co_* is the diameter of the associated colony. Experiments were conducted in triplicate.

### Exopolysaccharide (EPS) and biosurfactant screening

Putative biosurfactant-producing isolates were screened on TW80-MSM to assess for their cultivability hit rate using equation 1, whereas putative exopolysaccharide (EPS)-producing isolates were screened on CRA to evaluate for the occurrence of the characteristic black-pigment reaction. The number of isolates that formed black colonies on CRA was counted, and blackenability was determined with this equation:

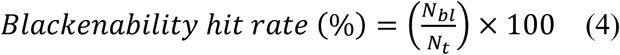

in which *N_bl_* is the number of black color-change indicated isolates after screening and *N_t_* is the number of total isolates before screening.

### Statistical analysis

All statistical analyses were performed using Statistical Package for the Social Sciences (SPSS) (Version 27, IBM, US). Normality was assessed, and analysis of variance (ANOVA) with Tukey post hoc test was used to determine significant differences in multiple comparisons. Paired T-test was conducted to determine the significance of clearability between non-selective and selective isolation groups.

## RESULTS AND DISCUSSION

### Mesophilic PLA-degrading bacteria were prevalent in the environment

The well-established industrial compostability of PLA suggested that its biodegradability would be present in the environment, and this idea was further strengthened by our development of a trained compost with mesophilic PLA degradability (14). To investigate these activities and delineate how our compost overcame the high-temperature requirement for PLA breakdown, we examined whether viable mesophilic PLA-degrading bacteria could be isolated from our trained compost and from a variety of environmental samples. The use of cultivable isolates likely resulted in many strains being missed, but identical procedures were applied across all samples to limit comparative bias. Furthermore, given that PLA biodegradation is not fully understood, we prioritized functional assays on viable bacteria over alternative strategies, such as genomics-based predictions, to assess biodegradation potential across samples.

A non-selective isolation strategy was conducted where microbial suspensions from different sites, including marine sites, were recovered and purified on NA, and distinct colonies were subsequently re-plated onto minimal agar with PLA as the sole carbon source (PLA-MSM, Fig. 1A). Phenotypic distinctions of isolates were validated by 16S rRNA sequencing, and all the isolates examined yielded the same designations by both methods (Table 1). In one case, two phenotypically distinct isolates (C-3-7, P-3-2) shared 97.86% 16S identity but were still classified as distinct, since different species can share >99% 16S identity (24, 25).

**FIG 1.**
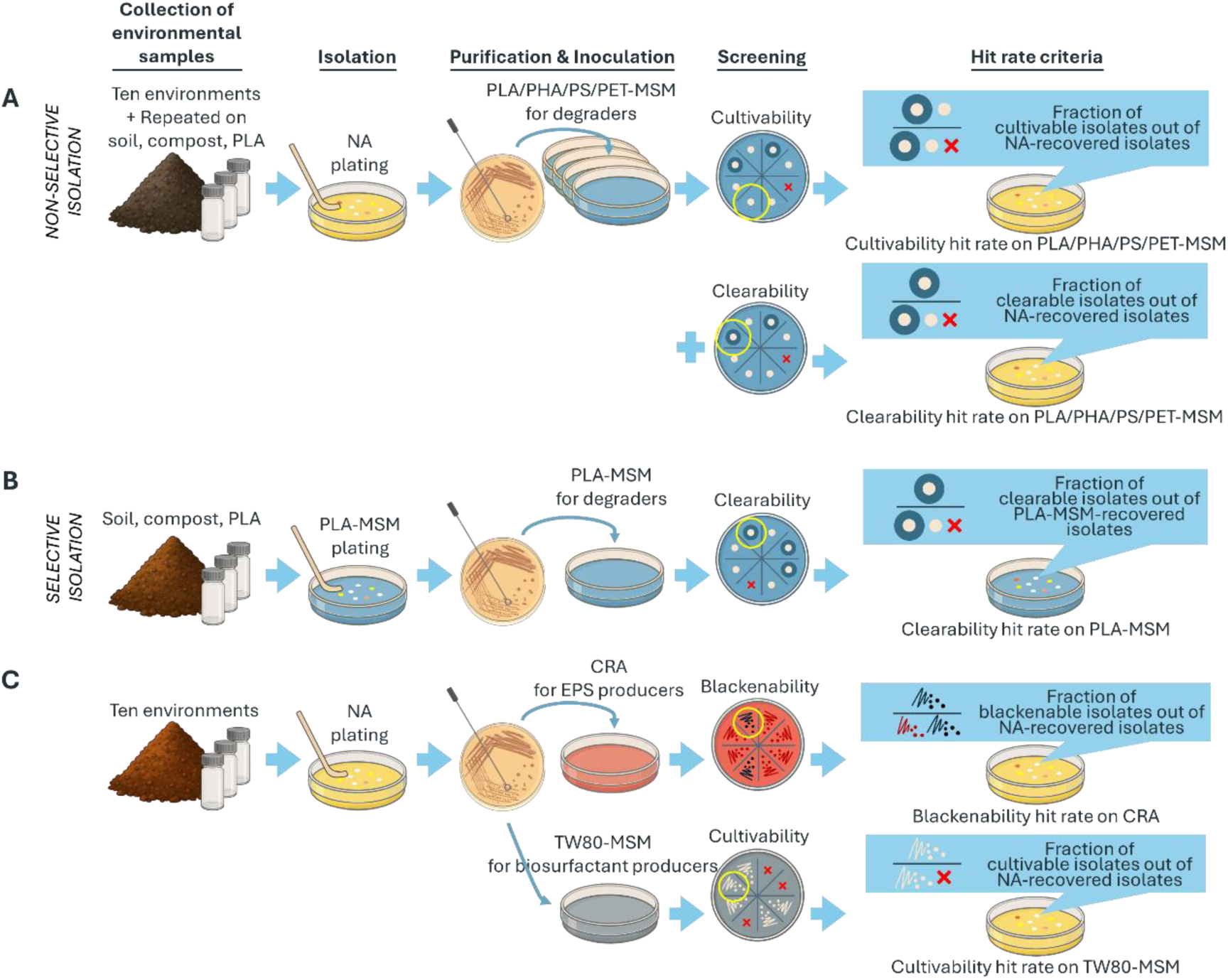
The experimental design illustrates the isolation methods and hit rate calculation criteria. (A) Non-selective isolation method where clearability and cultivability hit rates were determined from screening on PLA-, PHA-, PS-, and PET-MSM screening media out of the total isolates recovered on NA non-selective media. (B) Selective isolation method where clearability hit rate was determined from screening on PLA-MSM screening media out of the total isolates recovered on PLA-MSM selective media. Only soil, compost, and PLA environmental samples were recollected and proceeded using this method, and repeated in tandem with non-selective isolation for comparison. (C) Isolation of biosurfactant- and EPS-producing bacteria was recovered using non-selective isolation, where blackenability hit rate and cultivability hit rate were determined from screening on CRA for putative EPS producers and TW80-MSM for putative biosurfactant producers, respectively.

**Table 1.** Identity analysis of a subset of PLA-clearing isolates via 16S rRNA sequencing.

| Isolate groups <sup>a</sup> | Putative taxon based on 16S rRNA <sup>b</sup> | Pairwise comparison | Phenotypic determination | Designation by 16S rRNA | Pairwise 16S similarity (%) <sup>c</sup> |
| --- | --- | --- | --- | --- | --- |
| Chalky appearance |  |  |  |  |  |
| C-1-3 | *Streptomyces* sp. | C-1-3 and C-3-7 | Distinct | Distinct | 91.74 |
| C-3-7 | *Streptomyces* sp. | C-3-7 and P-3-2 | Distinct | Distinct | 97.86 |
| P-3-2 | *Streptomyces* sp. | P-3-2 and C-1-3 | Distinct | Distinct | 89.43 |
| Umbonate and translucent appearance |  |  |  |  |  |
| C-1-2 | *Paracidovorax* sp. | C-1-2 and C-2-2 | Distinct | Distinct | 92.91 |
| C-2-2 | *Comamonas* sp. | C-2-2 and C-1-8 | Distinct | Distinct | 86.25 |
| C-1-8 | *Achromobacter* sp. | C-1-8 and C-1-2 | Distinct | Distinct | 85.26 |
| Yellow and opaque appearance |  |  |  |  |  |
| C-3-1 | *Cellulosimicrobium* sp. | C-3-1 and C-7-3 | Distinct | Distinct | - <sup>d</sup> |
| C-7-3 | *Flavobacterium* sp. | C-7-3 and P-1-5 | Distinct | Distinct | 82.14 |
| P-1-5 | *Diaphorobacter* sp. | P-1-5 and C-3-1 | Distinct | Distinct | - <sup>d</sup> |
| Irregular and glossy appearance |  |  |  |  |  |
| C-3-3 | *Cellulosimicrobium* sp. | C-3-3 and C-3-5 | Distinct | Distinct | 81.99 |
| C-3-5 | *Leucobacter* sp. | C-3-5 and P-2-1 | Distinct | Distinct | 80.17 |
| P-2-1 | *Diaphorobacter* sp. | P-2-1 and C-3-3 | Distinct | Distinct | 82.35 |
| Irregular and shimmery appearance |  |  |  |  |  |
| C-2-9 | *Diaphorobacter* sp. | C-2-9 and S-2-6 | Distinct | Distinct | 93.33 |
| S-2-6 | *Diaphorobacter* sp. | S-2-6 and P-3-1 | Distinct | Distinct | - <sup>d</sup> |
| P-3-1 | *Pseudomonas* sp. | P-3-1 and C-2-9 | Distinct | Distinct | 80.98 |
| Lobate appearance (non-distinct test) |  |  |  |  |  |
| C-1-1 | *Diaphorobacter* sp. | C-1-1 and C-2-4 | Non-distinct | Non-distinct | 100.00 |
| C-2-4 | *Diaphorobacter* sp. | C-2-4 and S-1-9 | Non-distinct | Non-distinct | 100.00 |
| S-1-9 | *Diaphorobacter* sp. | S-1-9 and C-1-1 | Non-distinct | Non-distinct | 100.00 |

<sup>a</sup> Each set of three isolates represents a group selected for high similarity in colony morphology.
<sup>b</sup> Putative taxon of isolates compared to 16S rRNA databases determined by BLASTn.
<sup>c</sup> Pairwise isolate-to-isolate comparisons of 16S rRNA similarities using BLASTn for distinct isolates.
<sup>d</sup> No nucleotide similarity detected.

This experimental design enabled calculation of cultivability, defined as the number of bacterial isolates from a sample that grew on PLA-MSM relative to NA; this value represents the fraction that metabolizes or tolerates PLA, thereby estimating biodegradation potential across samples. Trained-compost samples (PLA, compost) and soil had the highest PLA-MSM cultivability hit rates (49.63–53.07%; Fig. 2A), whereas other environments were significantly lower but detectable (5.36–28.21%), indicating a modest prevalence of PLA-tolerant bacteria across all environments (Fig. 2A). Prior studies also reported soil as a comparatively diverse reservoir of PLA-degrading bacteria (26, 27).

**FIG 2.**
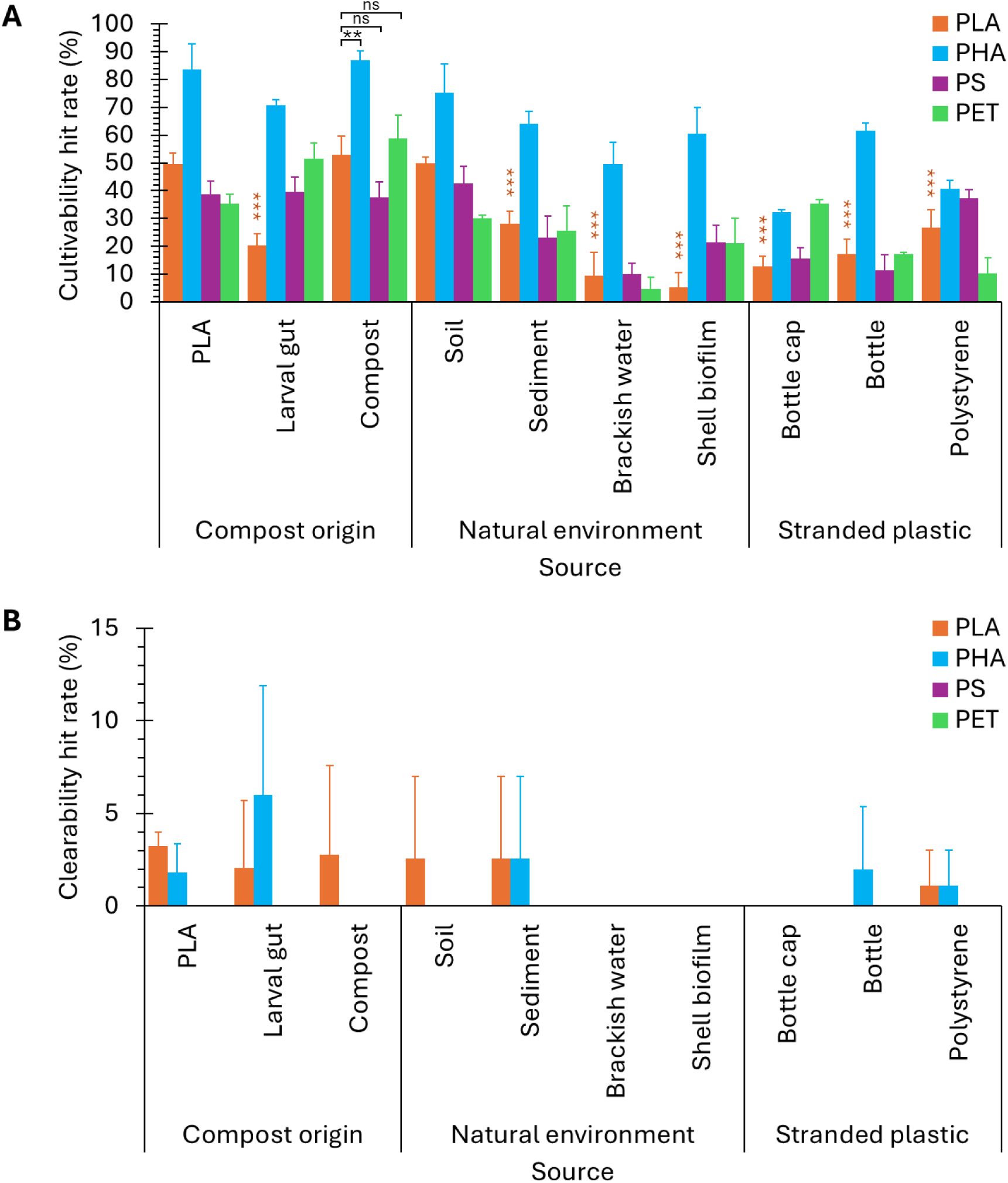
Comparisons of cultivability and clearability hit rates for different plastic types of various environmental samples. (A) Cultivability hit rates and (B) clearability hit rates of isolates from the ten environments screened on four MSM media supplemented with PLA, PHA, PS, or PET. Orange above-bar asterisks indicate statistical significance comparing the cultivability hit rates of the marked bars to that of the PLA sample, asterisks above brackets indicate comparisons between polymer groups (***: *P* < 0.001, **: *P* < 0.01, ns: *P* > 0.05, ANOVA with Tukey post-hoc test). Error bars indicate the standard deviation.

As growth alone does not indicate PLA hydrolysis, we further examined PLA clearability, which is a stricter proxy for hydrolysis (28). This was done by calculating clearability hit rates as percentages of isolates forming clearance zones on PLA-MSM (Fig. 1A). PLA-clearing isolates were detected (Fig. 2B), but were rarer than PLA-cultivable isolates (0-3.25%, Fig.2A), consistent with the greater stringency of the clearability criterion. While PLA clearability hit rates of microbes associated with the trained compost were among the highest (2.08-3.25% for PLA, larval gut, and compost), they were comparable to those of soil and marine sediment (both 2.56%, Fig. 2B). These results indicate that PLA clearability is broadly distributed in the environment and not enriched in our trained compost.

The general environmental existence of PLA clearance abilities led us to interrogate whether similar activities also exist in the environment for another biodegradable bioplastic, PHA, via the same non-selective isolation strategy, and we included PET and PS as low-biodegradability petroleum-based plastics for comparisons (Fig. 1A). All environments showed cultivability on PHA, PET, and PS, with PHA as the highest and exceeding those of PLA (32.38–86.94%; Fig. 2A), possibly due to PHA being a biologically familiar, microbial intracellular carbon reserve converted into metabolic energy under scarce nutrient environments (29). Under the stricter clearability criterion, no isolates cleared PET or PS (Fig. 2B). PHA clearability was much lower than PHA cultivability but occurred at rates similar to PLA (compare Fig. 2B to Fig. 2A), supporting that PHA degradability is also environmentally common (30) (https://doi.org/10.1007/s13762-024-05831-1). Overall, PLA- and PHA-clearing activities occured in a small subset of bacteria in many environments, whereas PET and PS biodegradability was much rarer. Interestingly, the limited enrichment for PLA clearers in our trained compost suggested that mesophilic compostability could be rooted in or encompassing activities beyond PLA depolymerization (31) (https://doi.org/10.1007/s00203-023-03701-x).

### Selective isolation facilitated identification of PLA-clearing isolates

NA is a general, non-selective medium, thus screening strains by first plating on NA may obscure unabundant, slow-growing, or metabolically specialized bacteria such as those that can metabolize PLA (32, 33). Therefore, we conducted a separate set of selective isolation by plating microbial filtrates directly on PLA-MSM to increase the chance of isolating PLA-clearing strains and investigate whether higher levels of enrichment of PLA degradability from our trained compost would be observed (Fig. 1B). The numbers of independent PLA degraders obtained were divided by the numbers of cultivable isolates to obtain hit rate values from this selective screening method.

The PLA clearability hit rates determined with this selective isolation method for microbes associated with the PLA, compost, and soil samples were between 6.74-16.98%, and the PLA sample did not exhibit higher clearability levels compared to the compost and soil samples (Fig. 3). Interestingly, clearability hit rates obtained with the selective isolation method were significantly higher than those derived with the non-selective screening method (Fig. 3A). The non-selective plating approach yielded no detectable PLA clearability hit rates in bacteria from soil, and only 1.59-2.12% in the PLA and compost samples (Fig. 3A-B). In contrast, PLA-MSM plating increased PLA clearability hit rates in soil to 10.47%, in compost to 16.98%, and in compost-derived PLA to 6.74% (Fig. 3A-B). Clearability hit rate of compost samples was the highest using the selective isolation method, consistent with previous reports that composts in general are enriched for hydrolytic activities (8, 12, 34, 35). In summary, the selective screening approach helped to uncover functional degraders that were otherwise overlooked by general cultivation; nevertheless, the findings further demonstrated that our trained compost exhibited at best a small enrichment in PLA degradation activity compared to similar environmental samples.

**FIG 3.**
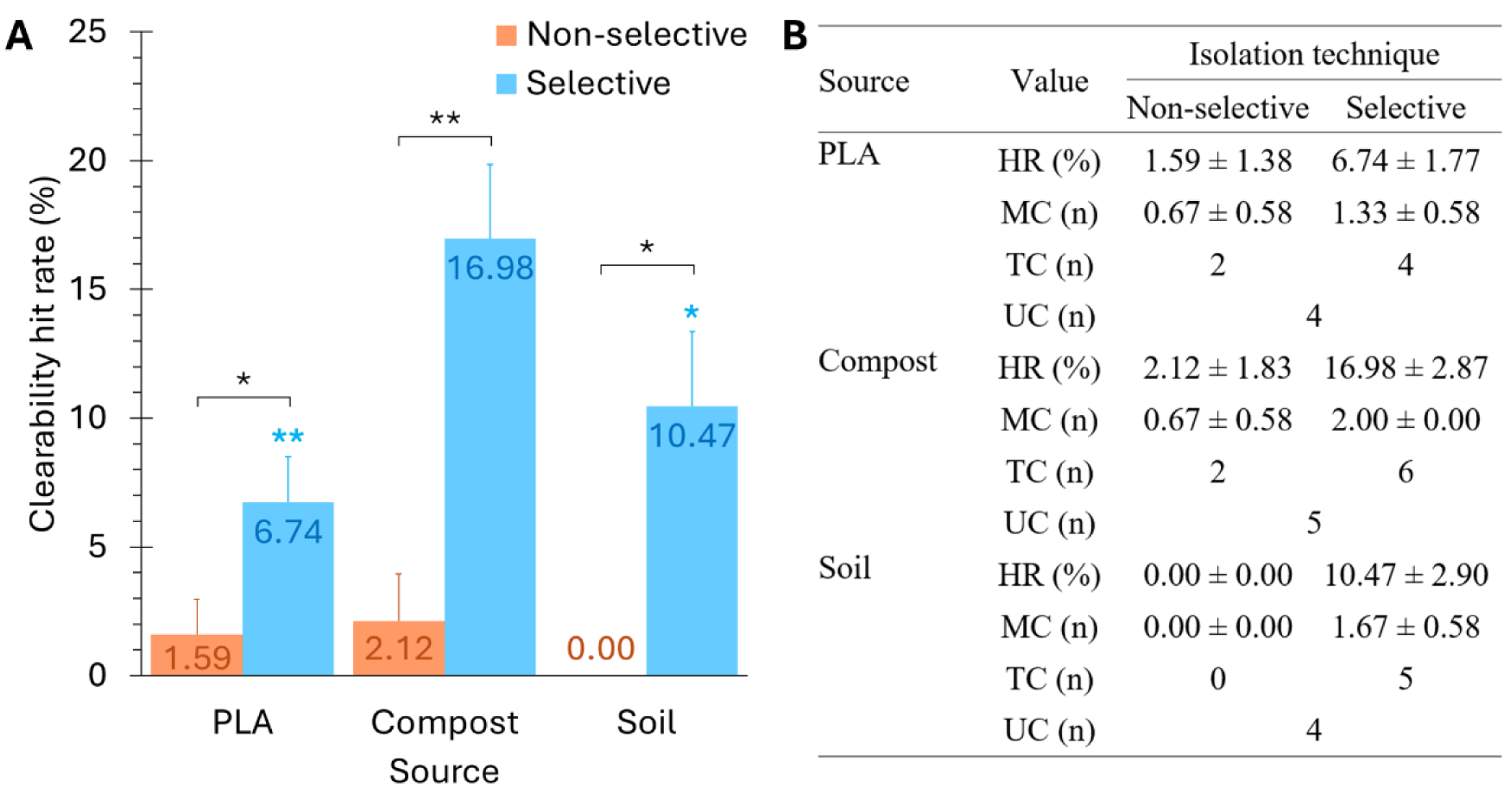
Selective isolation increased all PLA media clearability hit rates compared to non-selective isolation. (A) PLA media clearability hit rates of isolates from PLA, compost, and soil environments via non-selective and selective isolation. Blue asterisks above individual bars indicate selective isolation comparisons of PLA and soil to compost (ANOVA with Tukey post-hoc test), asterisks above brackets indicate comparisons between non-selective and selective isolation techniques (paired T-test, **: *P* < 0.01, *: *P* < 0.05). Error bars indicate the standard deviation. (B) Counts for the number of isolates (n) corresponding to the plotted clearability hit rates in (A). Hit rate (HR): percentage of clearance-causing isolates out of total recovered isolates on PLA-MSM screening (mean ± SD). Mean count (MC): mean number of isolates (mean ± SD). Total count (TC): pooled distinct isolates across triplicated experiments. Unique count (UC): distinct isolates after replicates were excluded.

### PLA and compost isolates showed increased PLA-clearing activity

Our trained compost did not appear to harbor a significantly higher fraction of PLA-clearing bacteria compared to several other environments. This led us to wonder if the compost has instead selected for strains that have higher PLA breakdown efficiencies (34, 36), which would help bring to light the elevated PLA biodegradability of our trained compost. This idea was tested by PLA clearing assays using emulsified PLA in PLA-MSM to ensure homogeneous polymer distribution and reduce clearance inconsistencies. All distinct PLA-clearing isolates derived from PLA, compost, and soil samples via both the non-selective and selection isolation methods were analyzed.

We compared isolates by measuring colony and clearance diameters and computing clearance indices (clearance diameter over colony diameter) to normalize for growth rate differences. This index prevented underestimation of slow-growing but active degraders and overestimation of fast-growing but weak degraders. We found that the trend for clearance efficiency was highest for PLA, followed by compost, and lowest for soil, with the average clearing efficiency for PLA being two-fold higher than that of soil. This implied that PLA-clearing isolates found on PLA surfaces were the most effective. For example, clearance index data across days 7, 14, and 21 revealed that isolate P-1-5 (PLA-derived) exhibited the highest clearance index (2.25) as well as the largest clearance diameter (62.30 mm; Fig. 4A). The superior clearance index of isolate P-1-5 suggested its potential as a true PLA-degrading specialist with high-activity PLA depolymerizing ability (29, 37). However, additional evidence, such as gene presence and growth absence in no-carbon or inert-polymer controls, is required to confirm true PLA mineralization capacity.

**FIG 4.**
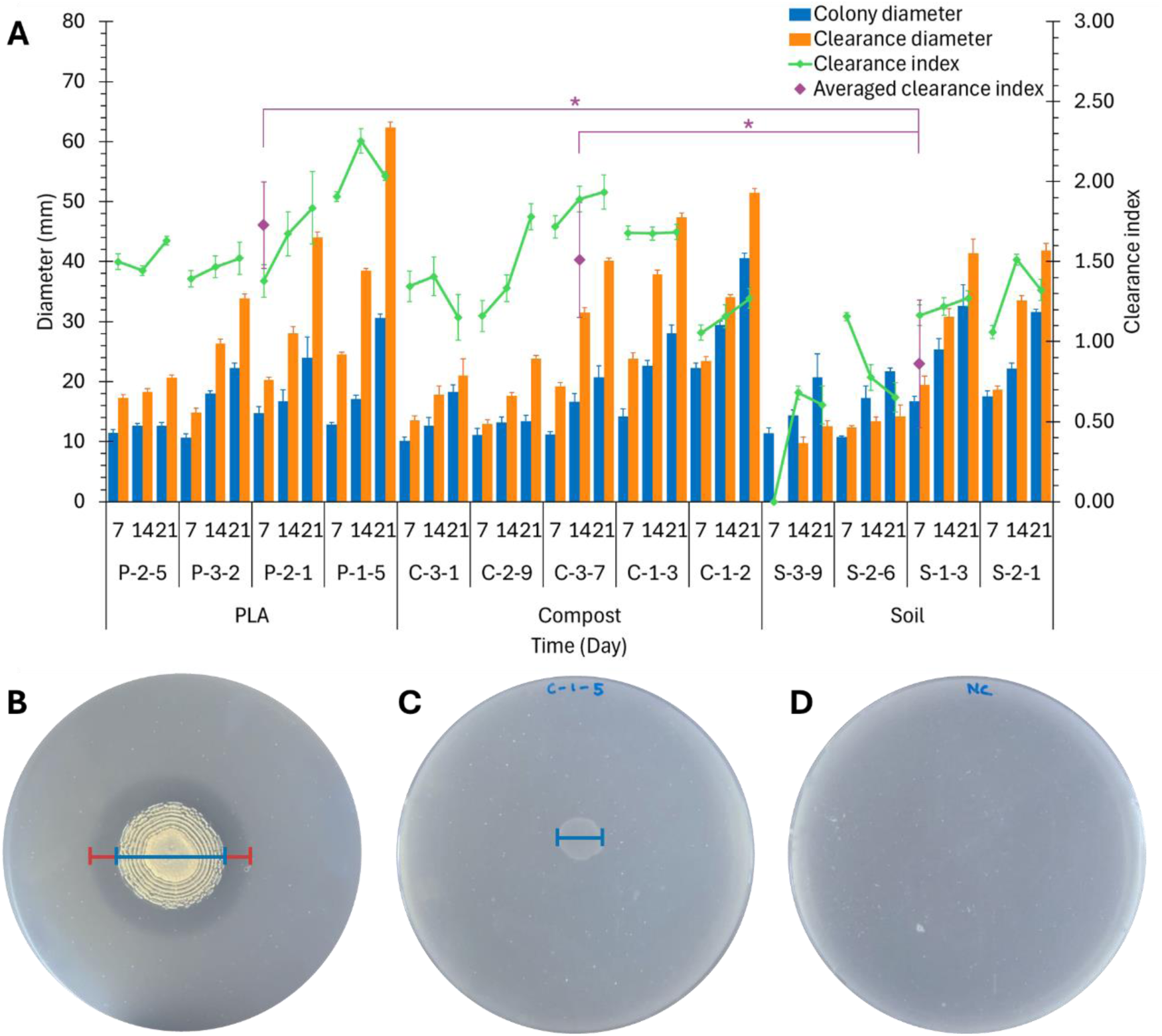
Quantitative analyses of PLA clearance for unique PLA-clearing isolates derived from PLA, compost, and soil. (A) Colony diameters, clearance diameters, and clearance indices of bacterial isolates from PLA, compost, and soil environmental samples incubated after 7, 14, and 21 days on PLA-MSM medium. Net clearance can be inferred from the gap between the colony and clearance diameters. Colony diameter, clearance diameter, clearance index, and averaged clearance index are indicated by blue bars, orange bars, green lines, and purple markers, respectively. Asterisks indicate averaged clearance index comparisons between environmental isolate groups (*: *P* < 0.05, ANOVA with Tukey post-hoc test). Error bars indicate the standard deviation. Representative photographs of 90-mm Petri dishes showing (B) halo presence by clearance-causing P-3-2 isolate, (C) halo absence by non-clearing C-1-5 isolate, and (D) uninoculated PLA-MSM as a negative control. Measurement lines of clearance diameter and colony diameter on dishes are indicated by the red and blue lines, respectively.

A compost isolate, C-3-7, achieved the second-highest clearance index (1.93) despite only the fifth-largest clearance diameter (40.10 mm) and low colony diameter (20.73 mm; Fig. 4A). This profile implied strong secretion of extracellular depolymerases coupled to slow cellular proliferation, consistent with low-biomass, high-activity polymer-degrading specialists that invest substantial metabolic energy into polymer-to-monomer bioconversion rather than rapid growth (38). In contrast, compost isolate C-1-2 produced the second-highest clearance diameter (51.48 mm), but its clearance index was relatively low (1.27; Fig. 4A), indicating a growth-focused degradative phenotype (colony diameter of 40.55 mm). Such a profile could be reflective of its microbial growth being supported by alternative substrates (e.g., agarolysis or indirect facilitation) or limited depolymerase expression, thus it is less likely to be a major contributor to PLA degradation (39). Overall, our quantitative analysis indicated that exposure to PLA selects or induces higher extracellular depolymerization capacity and enriches functional specialists, whereas soil communities with minimal contact to PLA harbor mainly weak degraders (40).

Our analysis showed that the PLA-clearing activity in our trained compost was about two-fold higher than that of soil, an unselected environmental sample (Fig. 4A). This finding illustrated how exposure to PLA would lead to enrichment of microbes that can mediate its breakdown, likely using the degradation products as an alternative source of energy and creating a survival advantage of isolates with higher PLA degradability. The higher clearing activity in PLA- and compost-derived bacteria inferred that the PLA biodegradability of our trained mesophilic compost stems more from efficiency than relative abundance of the culturable mesophilic PLA-degrading bacterial community (41, 42). However, this moderate enrichment is most likely not sufficient to elucidate the full extent of the increase in PLA compostability at mesophilic temperatures of our trained compost compared to standard compost piles (14), whose PLA compostability at ambient temperature is negligible. Therefore, we propose that synergistic PLA-microbe-environment dynamics, including cross-feeding, co-metabolism, and matrix effects, support in situ PLA turnover in compost (43). The key implication is that optimizing PLA biodegradation in mesophilic systems may require more than engineering of depolymerization specialists.

### Biosurfactant and EPS production were not enriched in trained compost-derived isolates

To examine additional bioactivities that may augment mesophilic PLA biodegradation in our compost, we screened isolates for two microbial functions implicated in facilitating plastic biodegradation: biosurfactant and EPS production (44, 45). Biosurfactants could increase PLA wettability of hydrophobic surfaces, while EPS could concentrate biodegradative enzymes, retain breakdown products for co-metabolism, and detach microfragments to increase reactive surface area (46, 47). To assess enrichment of these surface-modifying bacteria between the trained compost and other environmental samples, the proportion of isolates with the targeted activity was determined through functional screening (Fig. 1C). Biosurfactant potential was assessed by growth on TW80-MSM, and EPS potential by colony blackening on CRA. While these assays do not directly assay production of biosurfactants and biofilm, they are valid proxies that allow higher-throughput screening of isolate potential.

Hit rates for biosurfactant activity were found to range from 10% to 45% of isolates across diverse environments with no significant increase in the trained compost, indicating a broad ecological ubiquity of biosurfactant production rather than selection specific within our trained compost (Fig. 5). Such universality was consistent with previous studies, supporting the interpretation that biosurfactant producers in general are ecologically common (48–50). We next examined whether EPS producers were enriched in our trained compost, and found that the percentages of isolates that gave black colonies on CRA were similar in most of our samples (Fig. 5), suggesting that EPS production was also not enriched in the cultivable portion of compost bacteria. Interestingly, the percentages of EPS producing isolates were the highest in two polymer-associated samples from marine environments (bottle cap and PS, Fig. 5), inferring that EPS-producing strains could be selected for on the surfaces of waste marine plastics. This may have been due to microbial EPS facilitating adhesion and supporting biofilm structure (23, 51, 52). Collectively, the cultivation of bacteria from the environmental samples studied showed that EPS and biosurfactant production were not majorly enriched in our trained compost.

**FIG 5.**
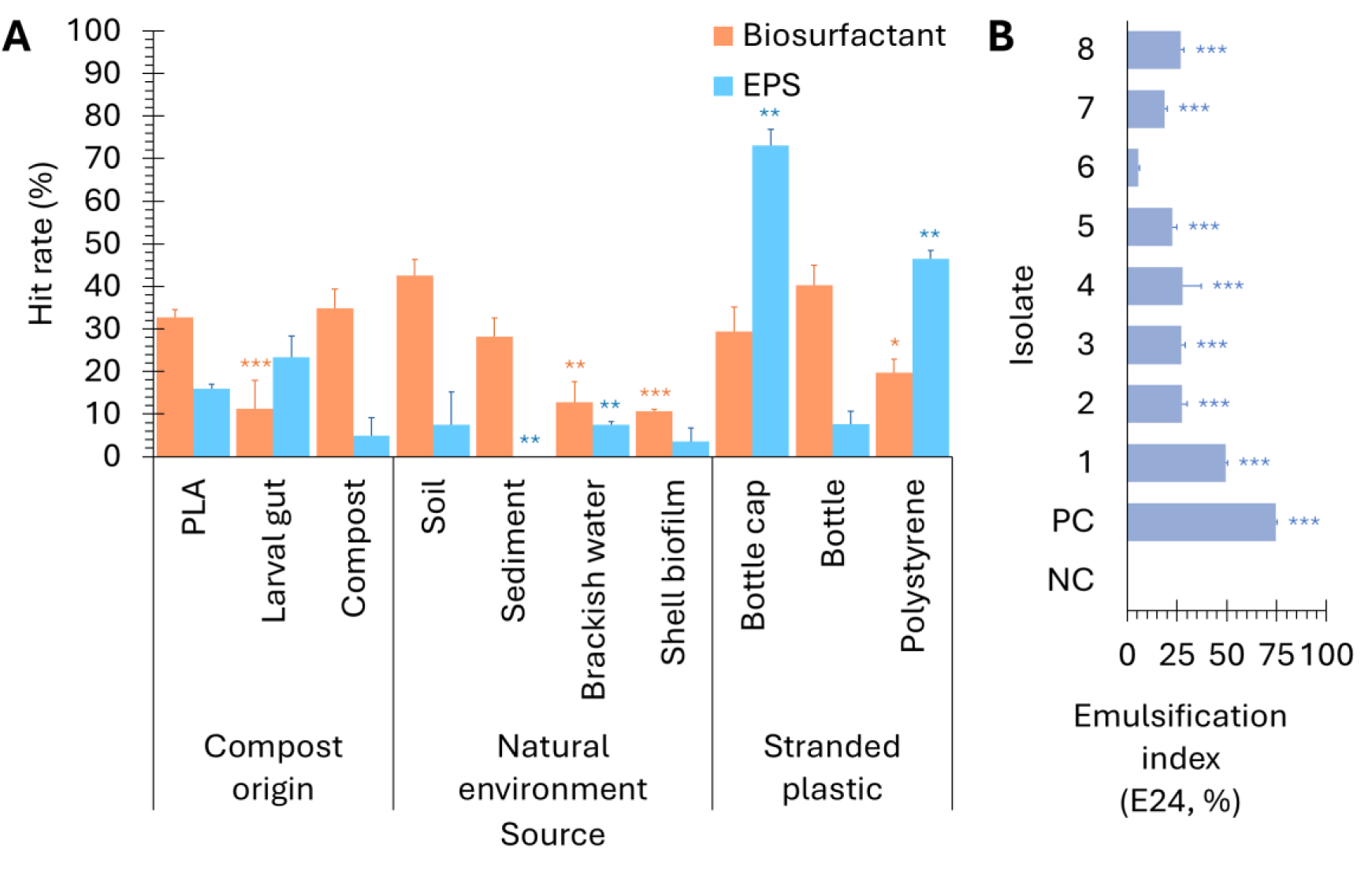
Mesophilic cultivable bacterial producers of biosurfactant and EPS are not enhanced in compost. (A) Hit rates of isolates tested positive for growth on TW80-MSM (biosurfactant-producing isolates, orange bars) and for producing black-colored colonies on CRA (EPS-secreting isolates, blue bars). Orange and blue asterisks above individual bars indicate comparisons to PLA samples (***: *P* < 0.001, **: *P* < 0.01, *: *P* < 0.05, ANOVA with Tukey post-hoc test). (B) Emulsification index (E24) tests on eight random isolates. TW80-MSM can be used as a proxy to rapidly screen for putative biosurfactant-producing bacteria (53). PC: 20% sodium dodecyl sulfate (SDS) solution. Blue-grey asterisks above bars indicate comparisons to negative control, uninoculated TW80-MSM (***: P < 0.001, ANOVA with Tukey post-hoc test). Values of standard deviation for each sample are indicated for each bar.

In summary, our survey revealed that our trained compost selected for higher efficiency PLA degraders compared to the general environment, providing a basis for why this compost could achieve mesophilic PLA compostability. The estimated enhancement of PLA compostability from this mechanism is estimated to be about two-fold, which is significant, but most likely not sufficient to explain the increase in PLA degradability in our trained compost compared to untrained compost. Taken together, our findings suggest that PLA biodegradability in our trained compost likely arose from a heterogeneous, multi-species consortium with diverse but complementary metabolic functions rather than from a single functional group.

## ACKNOWLEDGMENTS

The authors would like to thank the Yang lab for comments on the manuscript. This research was supported by an NSTC grant to S.Y.Y. (112-2311-B-182-003-MY3).

## AUTHOR CONTRIBUTIONS

Tan Suet May Amelia, Data curation, Formal analysis, Methodology, Visualization, Writing - original draft, Writing - review and editing | Shu Yuan Yang, Funding acquisition, Conceptualization, Methodology, Supervision, Writing - review and editing, Visualization

## REFERENCES

1. Rosenboom J-G, Langer R, Traverso G. 2022. Bioplastics for a circular economy. Nat Rev Mater 7:117–137.

2. Brizga J, Hubacek K, Feng K. 2020. The unintended side effects of bioplastics: Carbon, land, and water footprints. One Earth 3:45–53.

3. Zhang R, Jia S, Li J, Xu Y, Chen H, Zhang X. 2025. Techno-economic assessment of a closed-loop circular economy for polylactic acid. ACS Sustain Chem Eng 13:11226–11237.

4. Withana PA, Yuan X, Im D, Choi Y, Bank MS, Lin CSK, Hwang SY, Ok YS. 2025. Biodegradable plastics in soils: Sources, degradation, and effects. Environ Sci Process Impacts 27:3321–3343.

5. Mayekar PC, Auras R. 2024. Accelerating biodegradation: Enhancing poly(lactic acid) breakdown at mesophilic environmental conditions with biostimulants. Macromol Rapid Commun 45.

6. McKeown P, Jones MD. 2020. The chemical recycling of PLA: A review. Sustain Chem 1:1–22.

7. Ruggero F, Gori R, Lubello C. 2019. Methodologies to assess biodegradation of bioplastics during aerobic composting and anaerobic digestion: A review. Waste Manag Res 37:959–975.

8. Al Hosni AS, Pittman JK, Robson GD. 2019. Microbial degradation of four biodegradable polymers in soil and compost demonstrating polycaprolactone as an ideal compostable plastic. Waste Manag 97:105–114.

9. Mistry AN, Kachenchart B, Pinyakong O, Assavalapsakul W, Jitpraphai SM, Somwangthanaroj A, Luepromchai E. 2023. Bioaugmentation with a defined bacterial consortium: A key to degrade high molecular weight polylactic acid during traditional composting. Bioresour Technol 367:128237.

10. Figueiredo SN, de Oliveira Dias AR, Leitão LV, Folkersma R, Janssen K, Sampaio AAS, Almeida IO, Azevedo JB, de Carvalho LH, Torres FDPM, Barbosa R, Alves TS. 2025. Development of graphite-reinforced PLA / PBAT composite filaments. Polym Compos 46:13677–13689.

11. Hajighasemi M, Tchigvintsev A, Nocek B, Flick R, Popovic A, Hai T, Khusnutdinova AN, Brown G, Xu X, Cui H, Anstett J, Chernikova TN, Brüls T, Le Paslier D, Yakimov MM, Joachimiak A, Golyshina O V., Savchenko A, Golyshin PN, Edwards EA, Yakunin AF. 2018. Screening and characterization of novel polyesterases from environmental metagenomes with high hydrolytic activity against synthetic polyesters. Environ Sci Technol 52:12388–12401.

12. Mayekar PC, Limsukon W, Bher A, Auras R. 2023. Breaking it down: How thermoplastic starch enhances poly(lactic acid) biodegradation in compost - a comparative analysis of reactive blends. ACS Sustain Chem Eng 11:9729–9737.

13. Mayekar PC, Auras R. 2024. Speeding it up: Dual effects of biostimulants and iron on the biodegradation of poly(lactic acid) at mesophilic conditions. Environ Sci Process Impacts 26:530–539.

14. Hsueh SW, Kurniadi AC, Amelia TSM, Lee C-F, Fugmann SD, Yang SY. 2025. Mesophilic compostability of polylactic acid and the associated microbiome as revealed by metagenomics. J Hazard Mater Lett 6:100161.

15. Lamberti FM, Román-Ramírez LA, Wood J. 2020. Recycling of bioplastics: Routes and benefits. J Polym Environ 28:2551–2571.

16. Wang Y, Hu T, Zhang W, Lin J, Wang Z, Lyu S, Tong H. 2023. Biodegradation of polylactic acid by a mesophilic bacteria *Bacillus safensis*. Chemosphere 318:137991.

17. Wang Q, Zou X, Kang S, Wang Y, Li Z. 2024. Degradation of polylactic acid/polybutylene adipate-co-terephthalate blend by *Papiliotrema laurentii* S2P4P isolated from agricultural soils. Polym Degrad Stab 227:110855.

18. Xu Y-F, Wang M-Z, Wu Q-C, Zhou X, Ge X-W. 2016. Synthesis and morphology control of raspberry-like poly(ethylene terephthalate)/polyacrylonitrile microspheres. Chin Chem Lett 27:195–199.

19. Ben Ayed H, Jridi M, Maalej H, Nasri M, Hmidet N. 2014. Characterization and stability of biosurfactant produced by *Bacillus mojavensis* A21 and its application in enhancing solubility of hydrocarbon. J Chem Technol Biotechnol 89:1007–1014.

20. Lee J-S, Bae Y-M, Han A, Lee S-Y. 2016. Development of Congo red broth method for the detection of biofilm-forming or slime-producing *Staphylococcus* sp. LWT 73:707–714.

21. Gómez-Godínez LJ, Cisneros-Saguilán P, Toscano-Santiago DD, Santiago-López YE, Fonseca-Pérez SN, Ruiz-Rivas M, Aguirre-Noyola JL, García G. 2025. Cultivable and non-cultivable approach to bacteria from undisturbed soil with plant growth-promoting capacity. Microorganisms 13:909.

22. Viti C, Quaranta D, De Philippis R, Corti G, Agnelli A, Cuniglio R, Giovannetti L. 2008. Characterizing cultivable soil microbial communities from copper fungicide-amended olive orchard and vineyard soils. World J Microbiol Biotechnol 24:309–318.

23. Howard SA, Carr CM, Sbahtu HI, Onwukwe U, López MJ, Dobson ADW, McCarthy RR. 2023. Enrichment of native plastic-associated biofilm communities to enhance polyester degrading activity. Environ Microbiol 25:2698–2718.

24. Lin P-T, Wu Y-W. 2025. Highly-accurate prediction of colorectal cancer through low abundance uncultivated genomes recovered using metagenomic co-assembly and binning approach. BMC Cancer 25:1418.

25. Valido E, Bertolo A, Stoyanov J. 2025. Quantitative profiling of bacterial communities via full length 16S rRNA gene sequencing with internal controls: Optimization and validation across diverse human microbiomes. BMC Microbiol 25:710.

26. Liu H, Li M, Yu H, Shi L. 2025. Effects of conventional and biodegradable microplastics on soil physicochemical properties and microorganisms. Agric Res 10.1007/s40003-025-00906-y.

27. Saccà ML, Caputo F, Warren Raffa D, Suproniene S, Kadziene G, Skersiene A, Trinchera A. 2026. Relation of microbial functional genes to soil organic carbon and aggregate stability under different site conditions and soil management practices. Agric Ecosyst Environ 396:110012.

28. Urbanek AK, Rymowicz W, Strzelecki MC, Kociuba W, Franczak Ł, Mirończuk AM. 2017. Isolation and characterization of Arctic microorganisms decomposing bioplastics. AMB Express 7:148.

29. Pérez-García P, Sass K, Wongwattanarat S, Amann J, Feuerriegel G, Neumann T, Bäse N, Schmitz LS, Dierkes RF, Gurschke MF, Wypych A, Bounabi H, de Divitiis M, Vollstedt C, Streit WR. 2025. Microbial plastic degradation: Enzymes, pathways, challenges, and perspectives. Microbiol Mol Biol Rev 10.1128/mmbr.00087-24.

30. Abbas MI, Amelia TSM, Bhubalan K, Vigneswari S, Ramakrishna S, Amirul A- AA. 2025. Bioprospecting waste for polyhydroxyalkanoates production: Embracing low carbon bioeconomy. Int J Environ Sci Technol 22:2737–2756.

31. Saleem M, Yahya S, Razzak SA, Khawaja S, Ali A. 2023. Shotgun metagenomics and computational profiling of the plastisphere microbiome: Unveiling the potential of enzymatic production and plastic degradation. Arch Microbiol 205:359.

32. Kamagata Y. 2015. Keys to cultivating uncultured microbes: Elaborate enrichment strategies and resuscitation of dormant cells. Microbes Environ 30:289–290.

33. McCully AL, Loop Yao M, Brower KK, Fordyce PM, Spormann AM. 2023. Double emulsions as a high-throughput enrichment and isolation platform for slower-growing microbes. ISME Communications 3:47.

34. Kawashima N, Tokuda J, Yagi T, Takahashi K. 2022. Isolation of a *Nocardiopsis chromatogenes* strain that degrades PLA (polylactic acid) in pig waste-based compost. Arch Microbiol 204:599.

35. Cucina M, De Nisi P, Trombino L, Tambone F, Adani F. 2021. Degradation of bioplastics in organic waste by mesophilic anaerobic digestion, composting and soil incubation. Waste Manag 134:67–77.

36. Stojanovski G, Bawn M, Locks A, Ambrose-Dempster E, Ward JM, Jeffries JWE, Hailes HC. 2025. Functional enrichment and sequence-based discovery identify promiscuous and efficient poly lactic acid degrading enzymes. Environ Sci Technol 59:8602–8613.

37. Kim SH, Cho JY, Nara-Shin, Hwang JH, Kim HJ, Oh SJ, Kim HJ, Bhatia SK, Yun J, Lee S-H, Yang Y-H. 2023. Revealing the key gene involved in bioplastic degradation from superior bioplastic degrader Bacillus sp. JY35. Int J Biol Macromol 244:125298.

38. Cortes-Tolalpa L, Salles JF, van Elsas JD. 2017. Bacterial synergism in lignocellulose biomass degradation – complementary roles of degraders as influenced by complexity of the carbon source. Front Microbiol 8:1628.

39. Allison SD, Martiny JBH. 2008. Resistance, resilience, and redundancy in microbial communities. Proc Natl Acad Sci 105:11512–11519.

40. Meyer Cifuentes IE, Degenhardt J, Neumann-Schaal M, Jehmlich N, Ngugi DK, Öztürk B. 2023. Comparative biodegradation analysis of three compostable polyesters by a marine microbial community. Appl Environ Microbiol 89: e01060–23.

41. Sun Y, Eckstein S, Niu X, Yermakov M, Grinshpun S, Song G, Sun G. 2024. Biobased triesters as plasticizers for improved mechanical and biodegradable performance of polylactic acid fibrous membranes as facemask materials. ACS Sustain Chem Eng 12:7964–7975.

42. Pathak VM, Navneet. 2017. Review on the current status of polymer degradation: A microbial approach. Bioresour Bioprocess 4:15.

43. Mu L, Ding J, Wang Y, Peng H, Tao J, Pulkkinen E, Si H, Zhang L, Li A, Li J. 2025. Anaerobic biodegradation of PLA at mesophilic and thermophilic temperatures: Methanation potential and associated microbial community. Environ Technol 46:2932–2944.

44. Soni N, Kumarasamy V, Gupta P, Singh SDK, Kamaraj C, Subramaniyan V, Selvan ST, Velu RK, Velramar B. 2025. Enhancement of low-density polyethylene biodegradation through the production of surface-active compounds by *Pluralibacter gergoviae* TYB1. Sci Rep 15:23270.

45. Muangchinda C, Naloka K, Pinyakong O. 2025. Enhanced soil biodegradation of low-density polyethylene (LDPE) by a synthetic bacterial consortium: Performance, mechanisms, and community dynamics. Sci Total Environ 996:180131.

46. Ge Z, Lu X. 2023. Impacts of extracellular polymeric substances on the behaviors of micro/nanoplastics in the water environment. Environ Poll 338:122691.

47. Ziervogel K, Kehoe S, De Jesus AZ, Saidi-Mehrabad A, Robertson M, Patterson A, Stubbins A. 2024. Microbial interactions with microplastics: Insights into the plastic carbon cycle in the ocean. Mar Chem 262:104395.

48. Chaida A, Chebbi A, Bensalah F, Franzetti A. 2021. Isolation and characterization of a novel rhamnolipid producer *Pseudomonas* sp. LGMS7 from a highly contaminated site in Ain El Arbaa region of Ain Temouchent, Algeria. 3 Biotech 11:200.

49. Oliveira EM de, Sales VHG, Andrade MS, Zilli JÉ, Borges WL, Souza TM de. 2021. Isolation and characterization of biosurfactant-producing bacteria from Amapaense Amazon soils. Int J Microbiol 2021:1–11.

50. Guo P, Xu W, Tang S, Cao B, Wei D, Zhang M, Lin J, Li W. 2022. Isolation and characterization of a biosurfactant producing strain *Planococcus* sp. XW-1 from the cold marine environment. Int J Environ Res Public Health 19:782.

51. Wright RJ, Erni-Cassola G, Zadjelovic V, Latva M, Christie-Oleza JA. 2020. Marine plastic debris: A new surface for microbial colonization. Environ Sci Technol 54:11657–11672.

52. Casillo A, Lanzetta R, Parrilli M, Corsaro MM. 2018. Exopolysaccharides from marine and marine extremophilic bacteria: Structures, properties, ecological roles and applications. Mar Drugs 16:69.

53. Qazi MA, Kanwal T, Jadoon M, Ahmed S, Fatima N. 2014. Isolation and characterization of a biosurfactant-producing *Fusarium* sp. BS-8 from oil contaminated soil. Biotechnol Prog 30:1065–1075.

